# Glycaemic variability underlies myocyte dysfunction and myocardial injury risk in diabetes

**DOI:** 10.1101/2025.01.29.635173

**Authors:** Yuanzhao Cao, Meredith A. Redd, Jennifer E. Outhwaite, Dalia Mizikovsky, Woo Jun Shim, Chen Fang, Zhixuan Wu, Dara Daygon, Terra Stark, Robin Palfreyman, Han Sheng Chiu, Clarissa Tan, Ulrich Thomas, Elena Dragicevic, Julian Sng, Helen Barrett, Emily Dorey, Sonia Shah, Kirsty R. Short, Nathan J. Palpant

**Affiliations:** Institute for Molecular Bioscience, The University of Queensland, Brisbane, QLD, Australia; Faculty of Medicine, School of Biomedical Sciences, The University of Queensland, Brisbane, QLD, Australia; Ochsner Clinical School-The University of Queensland School of Medicine, New Orleans, Louisiana, USA; Queensland Metabolomics and Proteomics Facility, Australian Institute for Bioengineering and Nanotechnology, The University of Queensland, Brisbane, QLD, Australia; Nanion Technologies GmbH, München, Germany; School of Chemistry and Molecular Biosciences, The University of Queensland, Brisbane, QLD, Australia; Royal Hospital for Women, Randwick NSW, Mater Research-The University of Queensland, UNSW Medicine, NSW, Australia; Mater Research Institute-The University of Queensland, South Brisbane, QLD, Australia; Australian Infectious Diseases Research Centre, The University of Queensland, Brisbane, QLD, Australia

**Author notes:** Co-corresponding authors **Professor Nathan Palpant, Associate Professor Kirsty Short**, The University of Queensland, Brisbane, Australia. Co-first authors (Yuanzhao Cao, Meredith A. Redd).

**Keywords:** Glycaemic Variability, Myocardial Infarction, Ischaemia-Reperfusion Injury, Human Induced Pluripotent Stem Cell, Cardiomyocytes, Co-morbidity, Contractility, Metabolism, Cardiac Cell Death, Mortality, UK Biobank Cohort

## Abstract

Heart disease is the leading cause of morbidity and mortality in individuals with diabetes, due largely to risks associated with ischaemic injuries such as myocardial infarction (MI). We use human population genetic data to demonstrate that current biomarkers of hyperglycaemia do not account for risk of post-MI mortality in diabetes patients. This study therefore systematically evaluates glycaemic stress underpinning cardiovascular risk in diabetes. Using *in vivo* and *in vitro* models, we demonstrate that glycaemic variability rather than hyperglycaemia alone is a dominant risk factor for heart muscle dysfunction and myocardial injury sensitivity in diabetes. These findings provide new preclinical models for mechanistic and drug discovery studies and inform strategies for managing cardiovascular outcomes in patients with diabetes.

## INTRODUCTION

Diabetes mellitus (DM) is a metabolic disorder characterised by chronic hyperglycaemia with the total number of patients predicted to rise to 643 million by 2030 and 783 million by 2045 ^1^. Heart disease is the most prevalent cause of mortality and morbidity in people living with diabetes, due largely to risks associated with ischaemic injuries such as MI ^2,3^. Individuals with diabetes have a higher likelihood of post-MI mortality ^4-6^. Remarkably, the cardiovascular death rate is 4.4-fold higher in diabetes alone without other traditional cardiovascular risk factors in comparison to people without diabetes in the same age group ^7^. This epidemiological data demonstrates that the co-morbidity of diabetes and MI is an increasingly significant public health issue ^8,9^.

Ischaemic injuries such as MI are caused by a profound reduction in oxygen supply to the heart. For metabolically active organs such as the heart ^10^, the loss of oxygen supply is life-threatening. To salvage tissue, cells rely on anaerobic glycolysis to produce energy in the absence of oxygen. While this method of fuel utilisation has potential short-term benefits, a failure to quickly re-establish oxygen supply leads to a build-up of lactic acid. During an MI, the absence of blood perfusion leads to accumulation of lactate which causes a dangerous acidic drop in tissue pH. The sustained drop in pH following tissue ischaemia causes irreversible cellular damage, which can lead to death, disability, and increased risk of subsequent fatal events. Dysregulated serum glucose levels in people with diabetes result in metabolic dysfunction and oxidative damage that amplify the severity of myocardial damage during acute ischaemic events, potentially influencing both acute and long-term outcomes of MI ^11,12^.

Typically, in healthy individuals, blood glucose levels remain relatively stable throughout the day including small and short-lived postprandial peaks. In individuals living with diabetes, postprandial glucose spikes are more frequent and higher in magnitude, leading to drastic differences in the glycaemic variability in patients ^13^. Emerging evidence suggests glycaemic variability is a major risk factor for cardiovascular disease ^14^. Several clinical studies show that glycaemic variability increases the risk of MI and heart failure (HF) in people living with diabetes ^12,15^. In individuals with type 2 diabetes, mortality from cardiovascular disease was found to be associated with variability in fasting blood glucose levels ^15^.

This study uses biobank-scale human population data ^16^ to demonstrate that common risk factors of cardiovascular disease in patients living with diabetes, especially those associated with glucose regulation, do not effectively predict the risk of post-MI mortality. This study, therefore, evaluates the relationship and underlying mechanisms between glycaemic stress and acute cardiovascular injury and mortality risk in diabetes. We provide new mechanistic insights and benchmark *in vivo* and *in vitro* protocols for modelling cardiovascular risk in diabetes. Evidence from human genetic and modelling studies highlights the need to improve risk prediction of post-MI mortality in clinical diagnostics and therapeutic development to improve management and care for individuals living with diabetes.

## RESULTS

### Epidemiological analysis of cardiovascular risk in patients with diabetes

Epidemiological studies evaluating acute post-MI mortality in patients with diabetes mellitus (DM) vs non-diabetic patients (non-DM) demonstrate a consistent and significant increase in 30-day mortality rates (including in-hospital mortality) during acute MI in DM patients (**Figure S1A**). Among patients that do survive, post-MI heart function is not significantly different between DM and non-DM populations (**Figure S1B**). We aimed to understand the relationship between current clinical biomarkers of heart disease risk and diabetes as they predict post-MI risk and mortality in patients with DM. First, we investigated how clinical biomarker profiles affect the risk of MI outcomes in DM and non-DM in the UK Biobank Cohort (**Figure S1C**). We confirmed the increased incidence of both MI (**Figure 1A**) and MI mortality (**Figure 1B**) in DM individuals compared to non-DM individuals. Next, we used a published list of measured risk factors associated with post-MI mortality reported by Wohlfahrt et al ^11^ to evaluate the contribution of each risk factor to MI or post-MI mortality outcomes in DM and non-DM individuals. We evaluated the interaction between these common cardiovascular disease biomarkers and diabetes on the incidence and mortality of MI in the UK Biobank (**Figure 1C**). The results demonstrate that most biomarkers confer risk concordantly in DM and non-DM individuals for the incidence of MI (**Figure 1C, Figure S1D-F, Figure S1G-I**). However, all biomarkers fail to explain the increased risk of post-MI mortality observed in DM individuals (**Figure 1C**). Notably, haemoglobin A1c (HbA1c), a routine diagnostic marker of blood glucose, was effective in predicting MI incidence in both DM and non-DM individuals but was significantly less effective in predicting post-MI mortality in DM individuals than non-DM individuals (**Figure 1C**). Indeed, both DM and non-DM individuals with higher HbA1C levels were more likely to have an MI than those with lower HbA1C levels (**Figure 1D**). In contrast, there was no difference in post-MI survival time between DM individuals with high levels of HbA1c compared to DM individuals with lower levels of HbA1c (**Figure 1E**). Together, these findings show that classical cardiovascular disease risk biomarkers, including common measures of hyperglycaemia, fail to capture the increased post-MI mortality associated with DM, highlighting a significant area of need for new clinical biomarkers.

**Figure 1.**
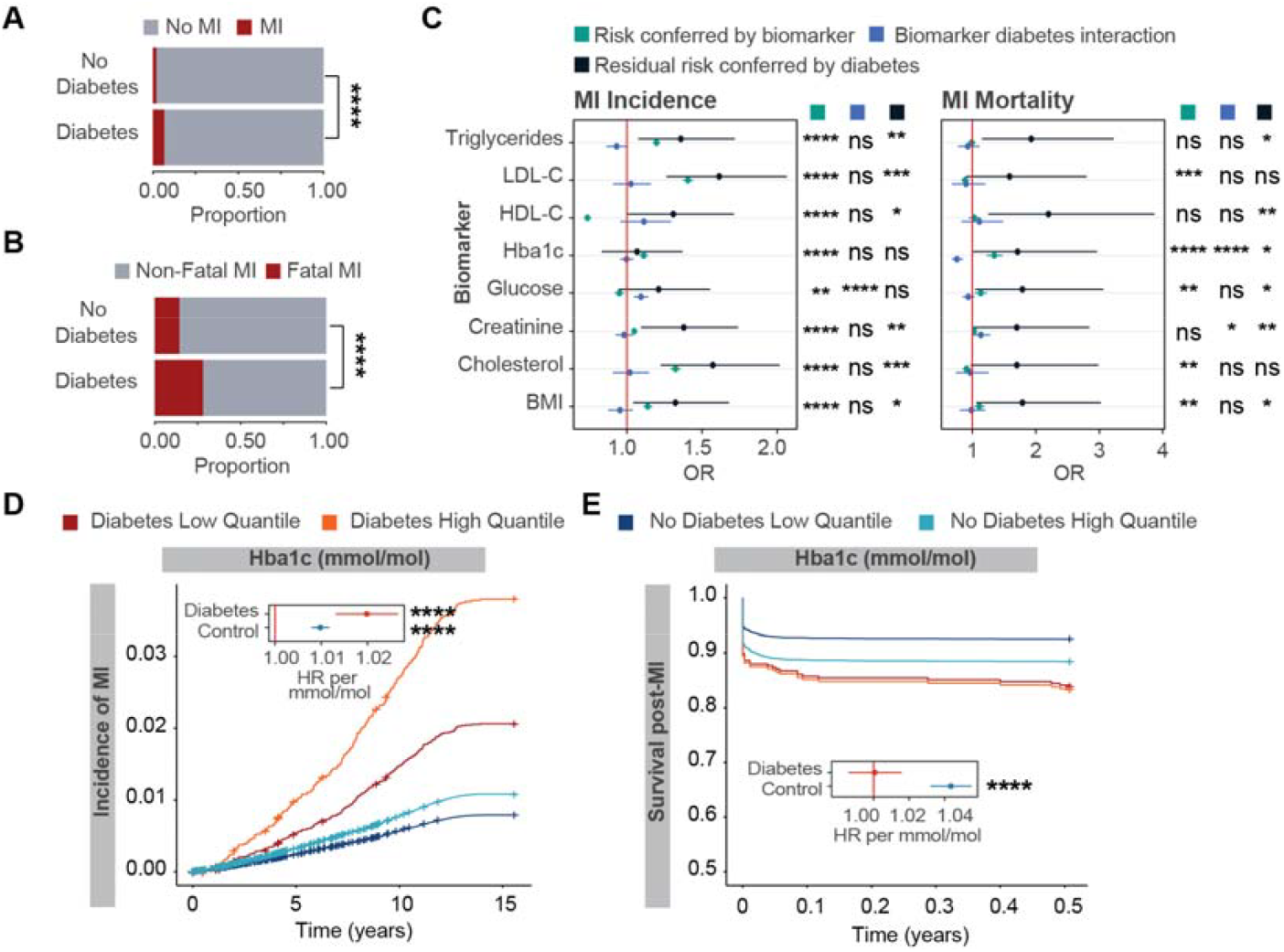
The risk of MI and mortality from MI in DM and non-DM individuals in the UK Biobank Cohort. (**A-B**). Incidence of MI (**A**) and mortality from MI (**B**) in individuals with and without diabetes. Chi-squared statistic used to determine significance. (**C**) Synergistic interaction between common cardiovascular disease biomarkers and diabetes on the incidence of MI and mortality in the UK Biobank. The predictor term (green dots) represents the risk conferred by the factor on MI incidence or mortality. The residual risk is defined as diabetes risk unexplained by the biomarker, and the term (black dots) represents the risk conferred by diabetes on MI incidence or mortality. The interaction term (blue dots) represents if the risk conferred by the risk factor is decreased or increased by the presence of diabetes. LDL, low-density lipoprotein; HDL, high-density lipoprotein; Hba1c, Hemoglobin A1C; BMI, body mass index. (**D-E**) Effects of known MI biomarker (Hba1c) on time to MI incidence (**D**) and mortality (**E**). Individuals stratified by diabetes diagnosis and further into the high quantile (top 30%) and bottom quantile (bottom 30%) of the biomarker levels. Data are presented as mean ± SEM and statistical analysis by Cox proportional hazards regression is used to determine hazard ratios and statistical significance (A-E). *P<0.05, **P<0.01, ***P<0.001, ***P<0.0001.

### *In vivo* modelling of glycaemic variability recapitulates the co-morbidity of diabetes and acute MI

Compared to healthy individuals with relatively stable blood glucose levels, individuals with diabetes have postprandial glucose spikes that are more frequent and higher in magnitude. Studies have demonstrated that drastic differences in glycaemic variability are a major risk factor for cardiovascular disease ^14^. We therefore evaluated the post-MI cardiovascular risk associated with constant hyperglycaemia vs glycaemic variability using an *in vivo* co-morbidity model involving a high-fat diet (HFD)-induced mouse model of prediabetes ^17^ exposed to an acute MI (**Figure 2A**). Mice were fed an HFD consisting of 40% calories from fat *ad libitum* for 18 weeks to induce a pre-diabetic state. An Alzet osmotic minipump was then implanted in the peritoneum. In variable glucose animals, the pump was loaded with PBS (released at a rate of 0.5 µL/h). These mice then received twice-daily intraperitoneal (i.p.) injections of glucose (10 mg), resulting in a variable glycaemic stress starting 1 day after surgery and continuing for 7 days (**Figure 2A**). Mice in the constant glucose group received a pump loaded with glucose (1.5 g/mL) and were administered twice daily injections of PBS for 7 days (**Figure 2A**). Thus, these two groups of mice received the same average amount of glucose over a 7-day period, one at a constant rate (cG) and the other in two boluses administered during the day (vG).

**Figure 2.**
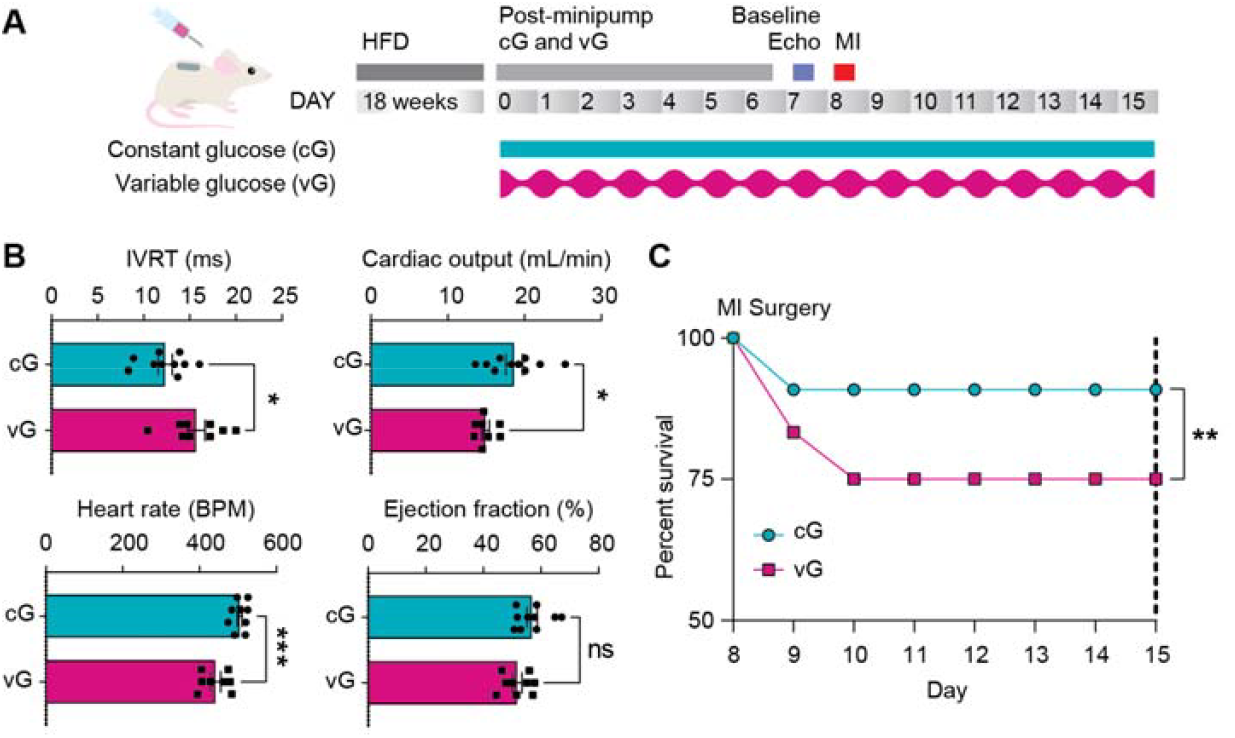
Comparative analysis of glycaemic stress for modelling ischaemia sensitivity in diabetes *in vivo*. (**A**) Schematic of protocol for an *in vivo* diabetic model of high fat diet (HFD)-induced diabetes exposed to variable (vG) or constant (cG) glycaemic stress followed by AMI surgery. (**B**) Echocardiographic analysis of heart function based on measures of isovolumic relaxation time (IVRT), cardiac output, heart rate, and ejection fraction. (**C**) Post-MI mortality analysis. For *in vivo* studies n=11-12 per group. Data are presented as mean ± SEM and statistical analysis by *t-*test (B) or Fisher’s exact test (C). *P<0.05, **P<0.01, ***P<0.001, ***P<0.0001.

On day 7, cG and vG animals were analysed for heart pump function by echocardiography. Both cG and vG animals showed equivalent contractile function based on analysis of ejection fraction (**Figure 2B**). vG animals showed a significant reduction in ventricular relaxation (prolonged isovolumic relaxation time) as well as reduced cardiac output, heart rate (**Figure 2B**). On day 8, animals were exposed to an acute MI by ligation of the anterior descending coronary artery for 40 minutes, followed by reperfusion ^18,19^. vG animals showed significantly greater acute mortality (25%) compared to cG animals (9.1%) within the first 48 hrs post-MI (**Figure 2C**). Collectively, these data demonstrate that glycaemic variability is a greater determinant than hyperglycaemia of the risk of acute mortality post-MI in diabetes.

### *In vitro* validation and mechanistic analysis of glycaemic variability in diabetic-induced myocyte dysfunction

Based on *in vivo* modelling results, we evaluated whether glycaemic variability influences cardiovascular function and disease risk *in vitro* using human pluripotent stem cell-derived cardiomyocytes (hPSC-CMs) ^20^. We tested hiPSC-CMs exposed to control media versus diabetic conditions (10 nM endothelin 1 (ET-1) and 1 μM cortisol) ^21^ involving either constant glycaemic stress (10 mM glucose, cG) or variable glycaemic stress (cyclic episodes of high (15 mM) and low (5 mM) glucose, vG) (**Figure 3A**).

**Figure 3.**
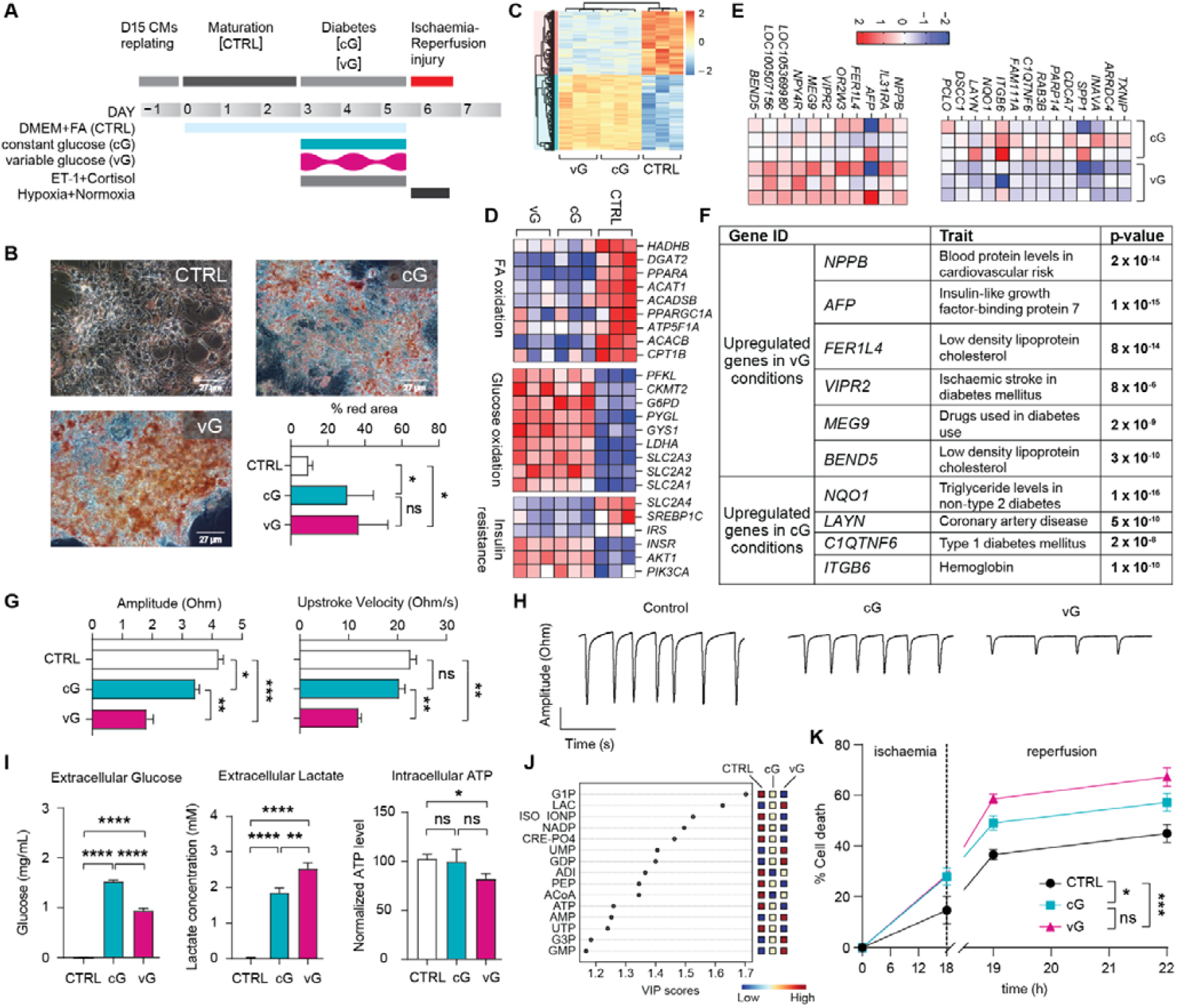
Comparative analysis of glycaemic stress for modelling ischaemia sensitivity in diabetes *in vitro*. (**A**) Protocol for hiPSC-CM diabetic and ischaemic stress model using a backbone media (Control) with diabetes-inducing neurohormonal mediators vs cells exposed to backbone + variable (vG) or constant glucose (cG). (**B**) Lipid uptake analysis based on Oil Red O staining (Scale Bar=27 µm). (**C-F**) RNA-seq of differentially expressed (DE) genes (**C**), diabetes-associated gene programs (**D**), and genes DE between cG and vG (**E**) with a table of gene-associated GWAS traits (**F**). (**G-H**) Contractility investigated by impedance-based analysis (CardioExcyte 96) showing bar plots (**G**) and raw contractility traces (**H**). (**I-J**) Metabolic analysis of glucose uptake, lactate secretion, and ATP production (**I**) with validation by unsupervised mass spectrometry (**J**). (**K**) Cell death measured by release of LDH in hiPSC-CMs exposed to *in vitro* ischaemia-reperfusion injury. For *in vitro* assays, n=3 biological replicates, each with 3 technical replicates. For metabolomics assay, n=3 biological replicates, each with 2 technical replicates. Data are presented as mean ± SEM and statistical analysis by One-Way ANOVA (B, G, I) or partial least squares - discriminant analysis (J) or Two-Way ANOVA (K). *P<0.05, **P<0.01, ***P<0.001, ***P<0.0001.

Compared to control, cG and vG cardiomyocytes showed significantly greater lipotoxicity characterised by increased intracellular uptake of lipid (Oil red O staining) (**Figure 3B**). RNA-seq analysis revealed that, compared to control, cG and vG conditions showed significant dysregulation of gene programs controlling insulin resistance, fatty acid (FA) oxidation, and glucose oxidation (**Figure 3C-D**). Notably, while there were few differentially expressed genes between cG and vG conditions, genes enriched in vG, including *NPPB* and *AFP*, have been significantly associatied with diabetes and cardiovascular disease (**Figure 3E-F**). We next evaluated contractile function using the impedance-based CardioExcyte system ^22^. These data show that cG and vG conditions both have impaired contractility compared to controls with vG conditions showing the greatest deficits in performance (**Figure 3G-H**), consistent with *in vivo* pump dysfunction under conditions of glycaemic variability (**Figure 2B**). Metabolic analysis of cG and vG conditioned hiPSC-CMs showed the greatest metabolic reprogramming in vG conditions based on increased glucose consumption, lactate production, and reduced ATP generation (**Figure 3I**), which we validated by liquid and gas chromatography mass spectrometry (**Figure 3J, Figure S2A-E**). Lastly, using an *in vitro* ischaemia-reperfusion (IR) injury model ^19^, we show that both glycaemic stress models had greater sensitivity to IR compared to control, however vG conditions showed the most significant sensitivity to ischaemia-induced cell death as measured by release of lactate dehydrogenase (LDH) (**Figure 3K**). These *in vitro* hPSC-CM perturbation studies mechanistically demonstrate that glycaemic variability is a significant risk factor for cardiomyocyte dysfunction and myocardial sensitivity to ischaemic stress.

### Blood serum from patients with high glycaemic variability induces contractile dysfunction in hiPSC-CMs

We next performed a proof-of-concept assay to test whether hPSC-CMs could reveal differences in cardiovascular function after exposure to plasma from patients with diabetes. We recruited diabetes patients (n=12) with glycaemic variability ranging from 22 to 41 (**Figure 4A-B**). Consistent with previous studies ^23^, we considered plasma measures >33 as high glycaemic variability. hPSC-CMs were exposed to donor serum samples added to standard backbone media (50:50 DMEM+FA:serum). These data show that plasma samples with high glycaemic variability cause a significant, acute reduction in contractility compared to plasma samples from patients with low glycaemic variability (**Figure 4C**). Notably, this difference in contractility was not observed if patients were stratified by their hyperglycaemic state (percent time very high), suggesting that glycaemic variability is a key determinant of cardiovascular outcomes (**Figure 4D**).

**Figure 4.**
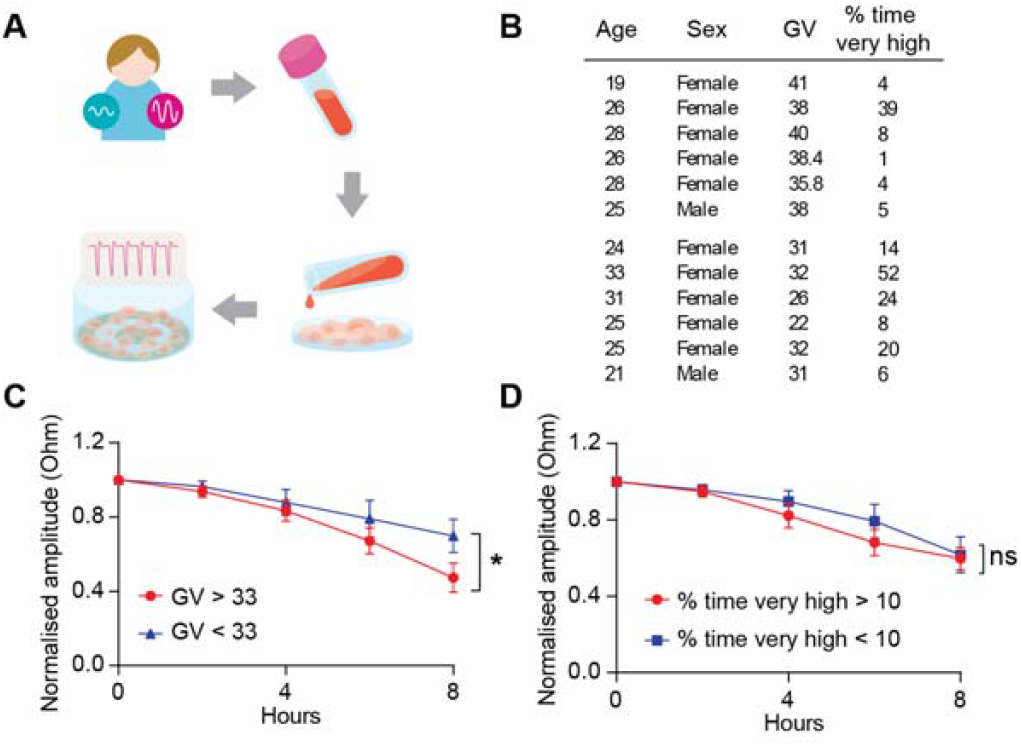
Clinical human blood samples evaluation on cardiac function. (**A-B**) Schematic of protocol (**A**) and patient details (**B**) for hPSC-CM contractility assays evaluating blood serum from patients with high vs low glycaemic variability. (**C-D**) Impedance-based contractility analysis comparing serum samples with high vs low glycaemic variability (**C**) or the percentage time very high (>13.9 mmol/L) (**D**). GV: glycaemic variability. For human serum samples assay, n=3 independent biological cardiomyocyte replicates, each with 1 technical replicate. Data are presented as mean ± SEM and statistical analysis by Two-Way ANOVA analysis (C-D). *P<0.05, **P<0.01, ***P<0.001, ***P<0.0001.

## DISCUSSION

This study evaluated mechanisms of glycaemic stress as a risk factor for cardiac ischaemic sensitivity and diabetes. We couple population genetics, clinical data, preclinical modelling and human blood serum studies to demonstrate a causal link between glycaemic variability as a risk factor in the pathophysiology of cardiac dysfunction in diabetes. A close alignment between epidemiological data and outcomes from *in vitro* and *in vivo* models provides compelling evidence for glycaemic variability as a culprit mechanism of cardiovascular disease risk in diabetes.

In contrast to markers reflecting consistently high blood glucose levels (i.e. elevated Hba1c), there is a growing body of evidence that glycaemic variability may play a significant role in susceptibility to cardiovascular disease ^14^. Clinical studies show that glycaemic variability increases the risk of MI and heart failure in people living with diabetes ^15^ and individuals with higher fasting blood glucose levels (>9.4 mmol/L), but not Hba1c levels, have a significantly higher risk of the most severe outcomes from MI ^11^. The current study uses cell, animal, and human population data to provide causal evidence that glycaemic variability directly impacts the health, function, and disease risk of cardiomyocytes in diabetes.

Furthermore, despite decades of research into mechanisms of ischaemic heart disease, there are no clinically approved drugs that block the acute injury response to cardiac ischaemia ^24^, making ischaemic heart disease the leading cause of death worldwide ^25^. While drug candidates for treating heart ischaemic injuries show promise in rodent models, none have managed to demonstrate efficacy in humans. At least in part, this is because most preclinical models do not recapitulate clinical patients who often have complex and co-morbid conditions like diabetes. Indeed, recent guidelines from the EU-Cardioprotection working group, IMPACT (IMproving Preclinical Assessment of Cardioprotective Therapies), recommend the inclusion of co-morbidity models to assess therapeutic efficacy of candidate drugs ^26^, including both cross-species and cross-platform modelling ^27^. While historical preclinical models ^21^ have used constant hyperglycaemia to induce diabetic cardiac stress, we demonstrate that glycaemic variability is a major variable in accurately modelling diabetes-associated cardiovascular risk. Taken together, the outcomes of this study provide new insights into disease mechanisms, establish new benchmark disease-relevant models to test new therapeutics and diagnostic assays, and provide new knowledge to improve management of cardiovascular health in patients with diabetes.

### Limitations of the study

The glucose dosages for the *in vivo* mouse model are based on a previously published mouse model of glycaemic variability ^17^. The 10-h blood glucose profile shows constant glucose dosage is 200 mg/dL, while the variable glucose dosage alternates between 200 and 400 mg/dL. These dosages were limited by the volume of the Alzet minipump, solubility of glucose, and efforts to match the total amount of glucose across the variable vs constant glycaemic stress models. Blood samples from patients with diabetes were not selected based on their medical history of MI, only on the basis of having constant glucose monitoring (CGM) data available. Based on existing ethics approvals, in this study we only had access to T1D patient serum samples and therefore performed a proof-of-concept study specifically assessing whether glycaemic variability impacts myocyte function. While T2D patients are at higher risk of heart disease, in Australia the National Diabetes Services Scheme (NDSS) provides federal funding to support CGM devices only for patients with T1D. This limited our ability to access blood samples with CGM data from T2D patients who are most relevant to our efforts modelling heart disease risk in diabetes. While data in this study demonstrate that hPSC-CM function is sensitive to patient-specific differences in glycaemic variability, future population scale retrospective cohort studies are needed to evaluate whether outcomes of hPSC-CM functional assays are predictive of cardiovascular disease outcomes in patients with diabetes.

## ACKNOWLEDGMENTS

This work has been supported by grant funding from the NHMRC (MRFCDDM000033 to NP and KS and 2007625 to NP as well as 2007919 and 1159959 to KRS), the Ian Potter Foundation (31111380 to NP), and the National Heart Foundation of Australia (106721 to NP).

