## Supplementary material for "Glycaemic variability underlies myocyte dysfunction and myocardial injury risk in diabetes": Methods

**Reagent or resource** **table**

| Reagent or resource | Source | Identifier |
| --- | --- | --- |
| Antibodies | | |
| Cardiac Troponin T | Thermo Fisher Scientific | Cat. #MA5-12960 |
| Phycoerythrin (PE)-conjugated sarcomeric α-actinin (SA) | Miltenyi Biotec Australia Pty | Cat. #130-123-773 |
| PE-conjugated anti-human isotype (IgG) | Miltenyi Biotec Australia Pty | Cat. #130-119-964 |
| Biological samples | | |
| Human serum samples | Hulme et al ^1^ | Mater Research Ethics ID (HREC/MML/55151 V2) and the University of Queensland Ethics ID (2019/HE002522). |
| Chemicals, peptides, and recombinant proteins | | |
| Vitronectin XF^TM^ | STEMCELL Technologies | Cat. #07180 |
| mTeSR™ Plus maintenance medium | STEMCELL Technologies | Cat. #100-0276 |
| Rock inhibitor Y-27632 (ROCKi) | STEMCELL Technologies | Cat. #72308 |
| Bovine serum albumin (BSA) | Sigma-Aldrich | Cat. #A9418-50G |
| L-Ascorbic acid 2-phosphate sesquimagnesium salt hydrate (ascorbic acid, AA) | Sigma-Aldrich | Cat. #A8960-5G |
| CHIR-99021 (CHIR) | STEMCELL Technologies | Cat. #72054 |
| XAV-939 | STEMCELL Technologies | Cat. #72674 |
| RPMI-1640 medium | Thermo Fisher Scientific | Cat. #11875093 |
| B-27 supplement plus insulin (50X, B27/insulin) | Thermo Fisher Scientific | Cat. #17504001 |
| Fetal bovine serum | ThermoFisher Scientific | Cat. #10099141 |
| 2.5% Trypsin | Thermo Fisher Scientific | Cat. #15090046 |
| Dimethyl sulfoxide | Sigma-Aldrich | Cat. #2650 |
| Saponin from quillaja bark | Sigma-Aldrich | Cat. #S7900 |
| 4% paraformaldehyde | Sigma-Aldrich | Cat. #158127-5G |
| DMEM no glucose medium | Thermo Fisher Scientific | Cat. #11966025 |
| MEM Non-essential amino acids | Thermo Fisher Scientific | Cat. #11140050 |
| L-Carnitine Hydrochloride | Merck Life Science Pty Ltd | Cat. #C0283-5G |
| Insulin-Transferrin-Selenium (100X) | Thermo Fisher Scientific | Cat. #41400045 |
| Linoleic Acid-Oleic Acid-Albumin (100X) | Merck Life Science Pty Ltd | Cat. #L9655-5ML |
| Taurine | Sigma-Aldrich | Cat. #T8691-25G |
| Creatine, anhydrous | Sigma-Aldrich | Cat. #C0780-50G |
| ENDOTHELIN 1 HUMAN PORCINE | Merck Life Science Pty Ltd | Cat. #E7764-50UG |
| HYDROCORTISONE | Merck Life Science Pty Ltd | Cat. #H4001-5G |
| Oil Red O | Sigma-Aldrich | Cat. #O0625-25G |
| Triton X-100 | Sigma-Aldrich | Cat. #X100 |
| D-(+)-GLUCOSE | Sigma-Aldrich | Cat. #G8270-100g |
| Versene Solution | Thermo Fisher Scientific | Cat. #15040066 |
| HEPES solution | Sigma-Aldrich | Cat. #H0887-100ML |
| Critical commercial assays | | |
| Colorimetric L-Lactate Assay Kit | Abcam | Cat. #ab65331 |
| Colorimetric HK Glucose Assay Kit | Sigma-Aldrich | Cat. #GAHK20-1KT |
| CellTiter-Glo Luminescent Assay | Promega | Cat. #G7571 |
| LDH cytotoxicity detection kit | Roche | Cat. #11644793001 |
| Deposited data | | |
| Bulk RNA-seq data | This manuscript | GEO: GSE279917 |
| Metabolomics data | This manuscript | MetaboLights: MTBLS11411 |
| Experimental models: Cell lines | | |
| Human iPSC: WTC-11 | Gift from Gladstone Institute  of Cardiovascular Disease, UCSF ^2^; ^3^ | RRID: CVCL_Y803 |
| Experimental models: Organisms/strains | | |
| Mouse: C57BL6/J male | Animal Resources Centre (Australia) | Ethics Committee ID: 2022/AE000203 |
| Software and algorithms | | |
| ImageJ | NIH | https://imagej.net/ |
| GraphPad Prism 9 (Version: 9.3.1) | GraphPad Software | https://www.graphpad.com/ |
| FACSDiva software | BD Biosciences | https://www.bdbiosciences.com/en-au/products/software/instrument-software/bd-facsdiva-software |
| FlowJo software (Version: 10.6.2) | Tree Star | https://www.flowjo.com/solutions/flowjo |
| DESeq2 | Love et al ^4^ | https://bioconductor.org/packages/release/bioc/html/DESeq2.html |
| CardioExcyte Control96 (Version: 1.4.7.1) | Nanion Technologies GmbH | https://www.nanion.de/products/cardioexcyte-96/ |
| Vevo LAB v5 software | VisualSonics | https://www.visualsonics.com/product/software/vevo-lab |
| MetaboAnalyst 6.0 platform | MetaboAnalyst | https://metaboanalyst.ca/ |
| Other | | |
| CardioExcyte 96 system | Nanion Technologies GmbH | https://www.nanion.de/products/cardioexcyte-96/ |
| CardioExcyte NSP-96 plates | Nanion Technologies GmbH | Cat. #201001 |
| ALZET osmotic minipumps | Alzet | Cat. #0000292 (ALZ) |
| Betadine solution (500mL) | Betadine | Cat. #12008-OR |
| Ethicon Vicryl Sutures, 6/0, 11mm, 3/8 Circle - J489G | EBOS healthcare | Cat. #JJ78849G |
| ETHICON 689G ETHILON NYLON SUTURES, 6/0 | EBOS healthcare | Cat. #JJ689G |
| Injectable glucose solution (GLUCOSE VIAL FOT 50% 50ML) | Clifford Hallam Healthcare (CH2) | Cat. #1383240 |
| 0.22um filter | Sigma-Aldrich | Cat. # CLs431218-50EA |
| RNeasy Mini Kit | QIAGEN | Cat. #74106 |

### **Resource availability**

### Lead contact

### Materials availability

This study did not generate new unique reagents.

### Data and code availability

Bulk RNA-seq has been deposited at the Gene Expression Omnibus database (GSE279917), the link is <https://www.ncbi.nlm.nih.gov/geo/query/acc.cgi?acc=GSE279917>, and the reviewer token is **itwxmacwrvmprsl**.

Metabolomics has been deposited at the MetaboLights database (MTBLS11411), the URL is www.ebi.ac.uk/metabolights/MTBLS11411, and the API token is **875697d3-abf9-4549-99ed-2d53ebb8bc34**.

This manuscript does not report any original code.

Any additional information required to reanalyse the data reported in this manuscript is available from the lead contact upon request.

**Generation and maintenance of human iPSC lines**

All human pluripotent stem cell studies were carried out in accordance with consent from The University of Queensland’s Institutional Human Research Ethics approval (HREC#: 2015001434). Cardiomyocytes generated in this study were derived from the WTC-11 hiPSC line (Gladstone Institute of Cardiovascular Disease, UCSF) ^2,3^. All WTC cells were maintained as previously described with slight adaptions ^5,6^. Briefly, iPSC cells were maintained in mTeSR Plus medium with supplement (Stem Cell Technologies, Cat.#100-0276) at 37°C with 5% CO_2_. Cells were cultured on Vitronectin XF (Stem Cell Technologies, Cat.#07180) coated plates (Nunc, Cat.#150318).

**Constant and variable glucose mice model**

All animal experiments were approved by the University of Queensland Animal Ethics Committee (2022/AE000203). Four-week-old C57BL6/J male mice were obtained from the Animal Resources Centre (Australia). Mice were placed on a high-fat diet (HFD) *ad libitum* consisting of 40% calories from fat (Specialty Feeds, Glen Forrest, Australia; SF04-001) for 18 weeks and supplied with water *ad libitum*.

To model glycaemic variability, mice were treated essentially as described previously ^7^. Mice receiving the HFD were intraperitoneally (i.p.) implanted with phosphate-buffered saline (PBS)-loaded 200 µL reservoir osmotic minipumps (Alzet) and administered 100 µL, 0.2 g/mL glucose i.p. twice daily for 7 days. Glucose solutions for the i.p injections were prepared by diluting two parts of sterile 50% w/v glucose solution (Phebra) in three parts of sterile PBS.

Mice with constant glucose levels received the HFD and Alzet osmotic minipumps loaded with 1.5 g/mL glucose (Sigma-Aldrich) were surgically implanted in the peritoneum. This pump releases a constant amount of glucose (1 μL/h). These mice also received twice daily i.p injections of PBS for 7 days. Accordingly, these two groups of mice received an equivalent glucose concentration every day for 7 days.

**Differentiation of iPSC-derived cardiomyocytes**

Cells were differentiated according to a modified monolayer platform protocol previously described ^5,6^. On day -1 of differentiation, hiPSCs were dissociated using Versene (Thermo Fisher Scientific, Cat.#15040066), plated onto Vitronectin XF-coated flasks (Nunc, Cat.#156367) at a density of 1.8 × 10^5^ cells/cm^2^, and cultured overnight in mTeSR Plus media supplemented with 10 µM Y-27362 ROCK inhibitor (Stem Cell Technologies, Cat.#72308). Differentiation was induced on day 0 by first washing cells with PBS, then changing the culture media to RPMI-1640 Medium (Thermo Fisher Scientific, Cat.#11875093) containing 3 μM CHIR-99021 (Stem Cell Technologies, Cat.#72054), 500 μg/mL bovine serum albumin (BSA, Sigma-Aldrich, Cat.#A9418), and 213 μg/mL ascorbic acid (AA, Sigma-Aldrich, Cat.#A8960). After 3 days of culture, the media was replaced with RPMI-1640 containing 5 μM XAV-939 (Stem Cell Technologies, Cat.#72674), 500 μg/mL BSA, and 213 μg/mL AA. On day 5, the media was exchanged for RPMI-1640 containing 500 μg/mL BSA, and 213 μg/mL AA without any additional supplements. From day 7 and onward, the cultures were fed every other day with RPMI-1640 containing 2% B27 supplement plus insulin (Life Technologies Australia, Cat.#17504001). Spontaneous beating was typically observed between days 9 and 11 of differentiation.

***In vitro* diabetic stress models with hiPSC-CMs**

Differentiated cardiomyocytes on day 15 were harvested using 0.5% Trypsin (Dilute 2.5% Trypsin stock 1:5 in Versene) with 1:1000 ROCKi). We plated 40,000 viable cells per well on Vitronectin XF-coated 96-well plate (Nunc, Cat.#167008) for *in vitro* disease modelling, and cells were incubated at 37°C, 5% CO_2_ overnight in replating medium (RPMI-1640+1X B27+5% FBS +1:1000 ROCKi) and were refreshed with standard medium (RPMI-1640/B27/Insulin) for 48 hrs. After 48 hrs, standard medium was exchanged for maturation medium (DMEM+FA) for 72 hrs. DMEM+FA is Dulbecco’s modified Eagle’s medium no glucose (Thermo Fisher Scientific, Cat.#11966025), 10 mM HEPES (Sigma-Aldrich, Cat.# H0887-100ML), 2 mM L-carnitine, 5 mM creatine, 5 mM taurine, 1 mM nonessential amino acids (Thermo Fisher Scientific, Cat.#11140050), 1× insulin-transferrinselenium (Thermo Fisher Scientific, Cat.#41400045), and 1× linoleic-oleic acid (Merck Life Science Pty Ltd, Cat.#L9655-5ML). After 72 hrs, DMEM+FA was exchanged for cG (DMEM+FA plus 10 mM glucose, 10 nM h-endothelin-1, and 1 µM cortisol), vG (DMEM+FA plus 15 mM or 5 mM glucose, 10 nM h-endothelin-1, and 1 µM cortisol) or DMEM+FA vehicle control for a further 60 hrs. vG model was treated with variant diabetic media containing glucose at 15 mM or 5 mM every 12 hrs cyclic episodes.

**Flow cytometry**

At the time of replating on day 15 of differentiation, a subset of cells (approximately 1 × 10^6^) was set aside for flow cytometry analysis of cardiomyocyte purity of differentiated cell populations. Cells were fixed with 4% paraformaldehyde (Sigma-Aldrich, Cat.#158127-5G), permeabilised in 0.75% saponin (Sigma-Aldrich, Cat.#S7900), and labelled with Phycoerythrin (PE)-conjugated sarcomeric α-actinin (SA) antibody (Miltenyi Biotec Australia Pty, Cat.#130-123-773) or Cardiac Troponin T (Thermo Fisher Scientific, Cat.#MA5-12960) or PE-conjugated anti-human isotype (IgG) control (Miltenyi Biotec Australia Pty, Cat.#130-119-964). Stained samples were analysed using a FACS CANTO II (Becton Dickinson) system with FACSDiva software (BD Biosciences). Data analysis was performed using FlowJo software 10 (version 10.6.2), and cardiac populations were determined with population gating from corresponding isotype controls. For all experiments, only cell preparations with greater than 80% sarcomeric α-actinin-positive or Cardiac Troponin T-positive CMs were used.

**Lipid accumulation**

Differentiated cardiomyocytes were replated on day 15 for lipid ORO staining. CMs cultured in 96-well plates were washed once in PBS and fixed in 4% paraformaldehyde for 15 mins at room temperature. Following 3 further PBS washes, cells were permeabilised in 0.1% Tween-20 in PBS for 10 mins at room temperature. Intracellular lipid was stained with 1.5 mg/mL ORO (Sigma-Aldrich, Cat.# O0625-25G) dissolved in isopropanol for 10 min. After 3 washing steps with PBS, image acquisition was performed. The percentage of ORO-stained area per cell was quantified using Image-J software.

**Processing bulk RNA-seq data** **for gene quantification**

hiPSC-CMs cultured in backbone media conditions or underwent cG and vG disease modelling, and cells were harvested for RNA preparation and genome-wide RNA sequencing. RNA-seq samples with 3 biological replicates for each group were aligned to hg38 using STAR aligner ^8^ with the default option, yielding more than 294M uniquely mapped reads in total with an average of 92.84% mapping rate. Gene-level read counts were quantified using htseq-count ^9^ using the NCBI hg38 RefSeq gene annotation

**Identify differentially expressed genes**

Genes with count per million (CPM) of at least 1 in at least 2 samples were kept for further analysis. Read counts were normalised using ‘calcNormFactors’ function available in EdgeR package, with TMM method ^10^. Negative binomial generalised linear models were fit. Differentially expressed (DE) genes were identified by comparing all possible pair-wise combinations of the conditions (false discovery rate < 0.001).

**Clustering genes by expression dynamics between disease and control conditions**

We collated DE genes from all comparisons and identified 2 gene clusters using hierarchical clustering with Euclidean distance and complete-linkage clustering method; first cluster representing 1,482 genes expressed more highly in cG and vG samples compared to the control, second cluster representing 1,266 genes expressed more highly in control samples compared to the diabetic groups.

**Contractile performance and CardioExcyte 96 recordings**

Differentiated cardiomyocytes were replated on day 15 of differentiation for physiological maturation and disease modelling. We plated 50,000 viable cells per well on Vitronectin XF-coated 96-well CardioExcyte NSP-96 plates (Nanion Technologies GmbH, Cat.#201001). Contractile parameters of beating hiPSC-CMs monolayers were recorded using the CardioExcyte 96 system (Nanion Technologies GmbH). Contractility recordings performed with CardioExcyte 96 were obtained with 1 ms time resolution and 1 kHz sampling rate. All experiments were conducted under physiological conditions using an incubation chamber integrated in the system (37 °C, 5% CO_2_ and 80% humidity). The CardioExcyte NSP-96 plates utilise two gold electrodes on a rigid surface in each well to study physiological changes on contractility via continuous impedance monitoring of CMs exposed to maturation or stress conditions. The parameters for detection were amplitude and upstroke velocity.

**Metabolic dysfunction**

Lactate secreted into the culture medium from CMs exposed to control, cG, vG conditions were measured with colorimetric L-Lactate Assay Kit (Abcam, Cat.#ab65331) according to the manufacturer’s instructions. In brief, supernatant samples were prepared (1:300 dilution) at 50 μL/well with Lactate Assay Buffer in a 96-well plate (Corning, NY). The Lactate Standard was diluted to 1 mM (adding 10 μL of the Lactate Standard to 990 μL of Lactate Assay Buffer), and mixed well. Addition of 0, 2, 4, 6, 8, 10 μL into each well individually preceded adjusting the volume to 50 μL/well with Lactate Assay Buffer. Subsequently, a total 50 μL/well Reaction Mix was prepared containing 46 μL Lactate Assay Buffer, 2 μL Lactate Substract Mix, 2 μL Lactate Enzyme Mix. After mixing, 50 μL of the Reaction Mix was added to each well, including the Lactate Standard and supernatants. The reaction was incubated for 30 min at room temperature before measuring OD_450nm_ in a microplate reader (TECAN, Infinite 200Pro).

Glucose in the culture medium was examined with a colorimetric HK Glucose Assay Kit (Sigma-Aldrich, Cat.#GAHK20-1KT). A standard curve was prepared (10 μL/well) corresponding to 1 mg/mL, 0.5 mg/mL, 0.25 mg/mL, 0.125 mg/mL, 0.0625 mg/mL, 0.03125 mg/mL, 0.0156 mg/mL of glucose into a 96-well plate (Corning, NY). The supernatant samples were diluted with deionised water (1:10) at 10 μL volume per well, 90 μL of Reagent was added to each well containing the standards and samples. The plate was incubated for 15 min at room temperature, before measuring the absorbance at 340 nm using a microplate reader (TECAN, Infinite 200Pro).

Cellular and tissue ATP levels were measured using CellTiter-Glo Luminescent Assay (Promega, Cat.#G7571) according to the manufacturer instructions. Cell numbers were determined by counting cells using a hemocytometer. Intracellular ATP level was quantified using Luminescent and was normalised to DMEM+FA control cardiomyocytes.

**Metabolomics**

hiPSC-CMs induced for disease modelling to cG and vG cells and vehicle control cells were harvested to extract intracellular metabolites. Metabolomics analysis was performed in the Queensland Metabolomics and Proteomics Facility (Q-MAP). Approximately 1 million cells were quenched with 1 mL ice cold 50% methanol added with internal standards (500 nM azidothymidine, U-^13^C-sorbitol (1.6 µM) and L-Valine ^13^C_5_,^15^N (8.3 µM)). Cells were homogenised using an OMNI Bead Ruptor Elite (NE486LL/A) prechilled to -1°C. The extracts were aliquoted for analysis of central carbon metabolism (CCM) through Liquid Chromatography Mass Spectrometry (LC-MS) and Gas Chromatography Mass Spectrometry Human Metabolome Database (GC-MS HMDB) open profiling ^11^. The aliquot for CCM analysis was purified by biphasic separation with chloroform. The aqueous layer was evaporated to dryness and resuspended in 100 uL 2% acetonitrile solution with 5 uM succinate d-6 as injection internal standard. 5 µL was injected into a Shimadzu Nexera liquid chromatography coupled to a Shimadzu 8060 triple quadrupole mass spectrometer. Compound separation was achieved using a Phenomenex Gemini NX-C18 3 µm x 150 mm x 2 mm (Part No. 00F-4453-B0) column with SecurityGuard guard column (Phenomenex Part No. AJ0-8367). Mobile phase consisted of A: aqueous solution of 7.5 mM tributylamine (pH 4.95 with acetic acid) and B: acetonitrile. The mass spectrometer was operated on scheduled multiple reaction monitoring (sMRM) in negative ionisation mode. Transition ions for each compound were as previously published ^11^, authentic reference standards serially diluted 15 times from 100 uM to 6 nM was used to quantify the detected peak areas. Concentrations were normalised using internal standards (AZT and succinate d6) by automated peak integration in LabSolutions Insight LCMS v4 sp2.

For GC-MS, 60 µL of sample extracts were dried under vacuum (Eppendorf Concentrator Plus) in glass pulled point inserts. Samples were derivatised by methoxyamination by the addition of 25 µL methoxyamine hydrochloride (30 mg/mL in pyridine, 2 hrs, 37°C, 900 rpm), followed by trimethylsilylation with 25 µL BSTFA + 1% TMCS (30 min, 37°C, 900 rpm, followed by 1 h ambient temperature incubation.

GC-MS analysis was performed on a Shimadzu GC/MS-TQ8050 NX system. 1 µL of derivatised sample was injected into the GC inlet set at 280°C in 1:10 split mode, and chromatographically separated using an Agilent DB-5 ms capillary column (30 m × 0.25 mm × 1 µm) with helium at 1 mL/min. A 100°C starting temperature was held for 4 min, then ramped at 10°C/min to 320°C and held for 11 min. Compounds were fragmented by electron impact ionisation and analysed in MRM mode. A high-quality matrix was manually curated using the Shimadzu LabSolutions Insight GCMS program (v.3.7 SP3). Metabolomics statistical data analysis was performed using MetaboAnalyst 6.0 platform.

***In vitro* IRI model with hiPSC-CMs**

Differentiated cardiomyocytes were replated on day 15 of differentiation for *in vitro* ischaemia assays. We plated 40,000 viable cells per well on Vitronectin XF-coated 96-well plate. After replating, the cells were maintained for an additional 3 days in RPMI + B27/insulin, followed by 3 more days in DMEM+FA medium. Matured CMs were then exposed to DMEM+FA /cG/vG media for disease modelling. To prepare media for ischaemia/reperfusion injury, 10x HBSS without sodium bicarbonate (Sigma-Aldrich, Cat.# H6136) was diluted to 1x concentration in sterile tissue culture-grade water. Solutions were buffered with 12 mM HEPES (for pH 7.4 media, Sigma-Aldrich) and the pH adjusted accordingly with 1 M NaOH. The medium was sterile filtered with 0.22 μm syringe filters (Millipore). Unless otherwise noted, the replated cells were treated overnight (18 h) in HBSS with or without peptide under either normoxic (~18.5% O_2_; 5% CO_2_) or hypoxic (0.5% O_2_; 5% CO_2_) culture conditions. For reperfusion experiments, the medium was refreshed with HBSS pH 7.4 after overnight incubation and cultured for 1 h and 4 hrs under in normoxic conditions.

To assess cell death, the supernatant was collected, and LDH levels were measured using a cytotoxicity detection kit (Roche, Cat.#11644793001). For all cell culture experiments, percent cell death was calculated using low and high controls. For low control (LC), cardiomyocytes were cultured overnight in standard culture media (RPMI + B27/insulin). For high control (HC), cells were cultured in RPMI plus B27/insulin supplements containing 1% Triton X-100 (Sigma-Aldrich, Cat.#X100).

**Mouse MI/R Surgery**

MI/R surgeries were performed on constant and variable glucose mice. Briefly, mice were anesthetized with isoflurane (4% for initial induction then reduced to 1.5-2% to maintain anaesthetic plane) and mechanically ventilated via endotracheal intubation. A lateral thoracotomy was performed, and ischaemia-reperfusion injury was induced by occlusion of the left anterior descending (LAD) artery for 40 minutes with a small piece of PE10 tubing placed approximately 2 mm below the left atrial appendage, held in place with 7-0 prolene suture (Ethicon, Somerville, NJ, USA). At the end of the ischemic period, the suture and tubing were removed, the chest was closed with 5-0 prolene suture (Ethicon, Somerville, NJ, USA), and animals were provided analgesia by subcutaneous injection of 0.05 mg/kg buprenorphine, extubated and allowed to recover.

**Echocardiography**

Echocardiography was performed 1 day prior to MI/R surgery (baseline) using a Vevo3100 ultrasound system (VisualSonics, Toronto, Canada) with a 25–55 MHz transducer (MX550D) as previously described ^12^. All image acquisition and analysis were conducted by a blinded observer. For each animal, parameter/s were measured at least three times and averaged. During image acquisition, mice were maintained under general anaesthesia (4% isoflurane for initial induction then reduced to 1.5-2%) with body temperature monitored and maintained at 37.0 ± 0.5 °C. Two-dimensional B-mode images were collected in parasternal long-axis view and analysis was carried out using Vevo LAB v5 software (VisualSonics, Toronto, Canada). Reported parameters include Isovolumic Relaxation Time (IVRT), Heart Rate, Cardiac Output, Ejection Fraction.

**Human serum sample collection and iPSC modelling**

A cohort of 12 people with clinically diagnosed T1D who were routinely using continuous, or flash glucose monitoring (CGM) devices were recruited between 8/6/21 and 11/11/21 as previously described previously ^13^. Inclusion criteria were 18-60 years of age, not pregnant at the time of study, non-smokers of nicotine cigarettes, minimum diabetes mellitus duration of two years and no known immune disease requiring immunosuppressants. Blood, clinical data, and two weeks of prior CGM data from these participants were collected at the point of recruitment. Plasma was isolated by centrifugation. This study was approved by Mater Research Ethics Committee (HREC/MML/55151 V2) and the University of Queensland Ethics Committee (2019/HE002522).

Differentiated cardiomyocytes were replated on day 15 of differentiation as described above. Human plasma samples were diluted 50:50 in DMEM+FA media, added to the hPSC-CMs and allowed to equilibrate for 12 hrs. Contractile parameters of beating hiPSC-CMs monolayers were recorded using the CardioExcyte 96 system (Nanion Technologies GmbH) as described above. Serum samples for each patient were analysed across 3 independent biological cardiomyocyte replicates.

**UK Biobank cohort analysis**

The UK Biobank cohort was used to perform all analyses. Individuals were excluded from the cohort if they were of non-Caucasian ethnicity, had suffered an MI prior to recruitment into the UK Biobank, or had mortality from non-MI-related causes. Individuals in the UK Biobank were defined to have suffered an MI according to hospital diagnosis record ICD10 code I21. Cause of death was determined as the primary cause of death in the NHS death registries. Individuals were defined as having type 2 diabetes mellitus (DM) based on hospital diagnosis record ICD10 code E11. Further, only individuals who had received diagnosis prior to sample collection for the UK Biobank were retained (Field 130708). Individuals diagnosed with other forms of diabetes (ICD10 codes E10, E12, E13, E14) or self-reporting diabetes without accompanying hospital records, or those taking glucose medication were excluded from the control cohort.

A panel of biomarkers and risk factors were selected from Wohlfahrt et al ^14^. To establish if there was an interaction between the presence of diabetes and these factors on the risk of MI, or the risk of mortality from MI, a logistic model was fitted on each risk factor, diabetes, and a product variable between them. Continuous risk factors were scaled. Age, sex, medication status for glucose, lipid and blood pressure medications and Townsend deprivation index were used as covariates. Medication status was accounted for as a binary variable for each category based on UK Biobank fields 6153 and 20003. Logistic regression analyses were performed using the *glm* function in R.

To further investigate relevant biomarkers, we used Cox Proportional Hazards regression models on time from sample collection to MI and time from MI to death. Models were fitted separately in the diabetic and non-diabetic cohorts. For each biomarker, these models were first fitted for the unscaled value on a continuous scale with full covariates. Next, the individuals in the top and bottom 30% of the biomarker were defined.

**Statistics and reproducibility**

CMs on day 15 were used from 3 independent hiPSC differentiation. Each experiment group was performed with a minimum of three biological replicates. Statistical analysis was performed using GraphPad Prism 9 (version 9.3.1) software. To compare two normally distributed groups, a Student’s *t-test* was performed. For three or more groups and the assessment of one parameter, a One-Way ANOVA statistical test was used. A Fisher’s exact test was used to compare post-MI mortality values from cG and vG categories. With multiple parameters and three or more groups, a Two-Way ANOVA was used. Significance was determined as *p < 0.05, **p < 0.01, ***p < 0.001 and ****p < 0.0001. Data are presented as mean ± SEM.
