## Supplemental Figures for "Glycaemic variability underlies myocyte dysfunction and myocardial injury risk in diabetes"

**SUPPLEMENTAL FIGURES AND LEGENDS**

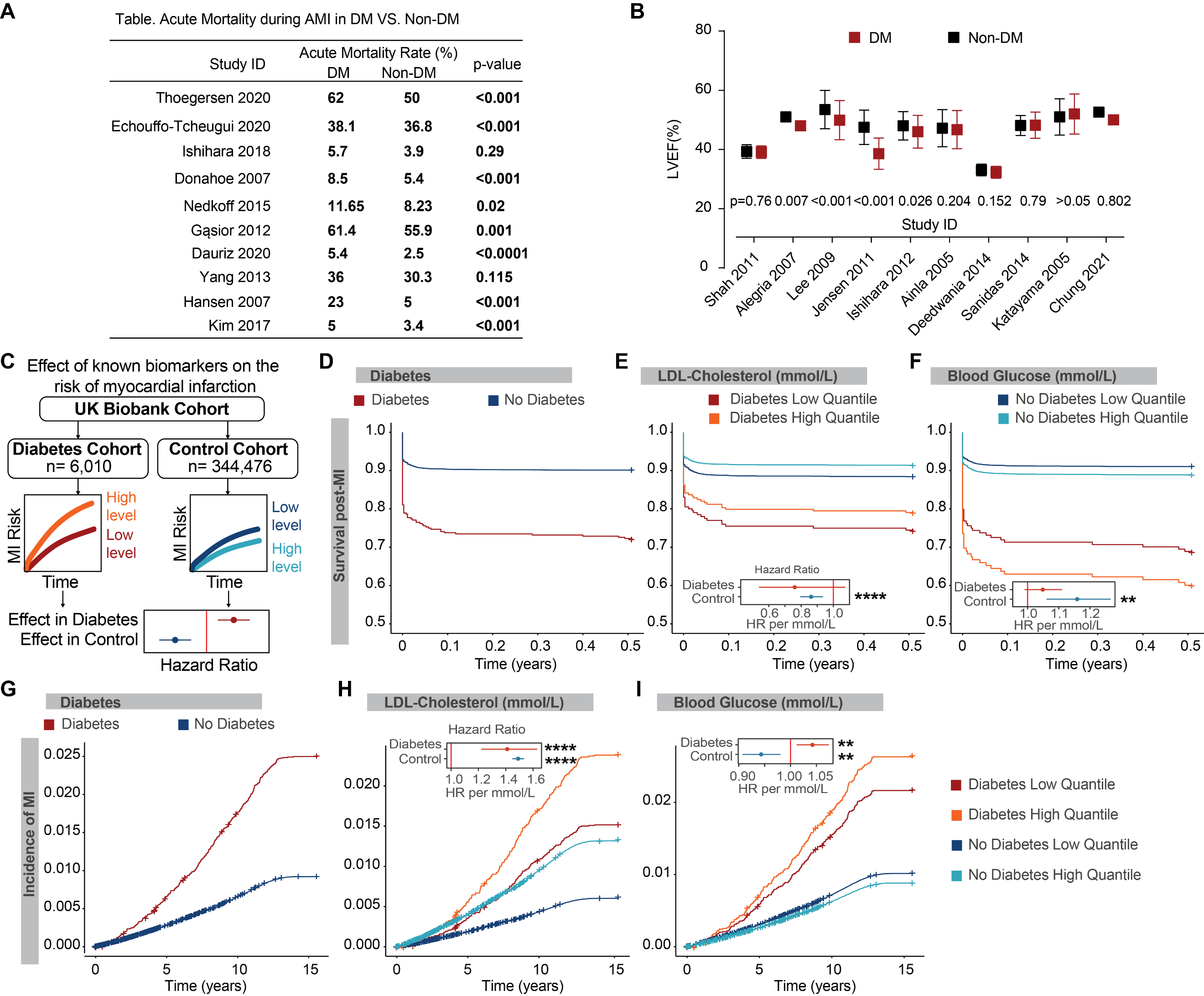

**Supplemental Figure 1. The risk of MI and mortality from MI in DM and non-DM individuals in clinical, epidemiological data and in the UK Biobank Cohort.** (**A**) Clinical and epidemiological summaries of acute mortality rates during AMI in DM and non-DM. (**B**) Clinical and epidemiological summaries of heart dysfunction on LVEF changes post-AMI in DM and non-DM. (**C**) Summary of analyses to characterise the association of blood biomarkers in individuals with and without diabetes to the incidence and mortality of MI in the UK Biobank. (**D**) Cox proportional hazards modelling time to mortality from recruitment to the UK Biobank following MI stratified by diabetes diagnosis. (**E-F**) Effects of known MI biomarkers on time to MI mortality. Individuals stratified by diabetes diagnosis and further into the high quantile (top 30%) and bottom quantile (bottom 30%) of the biomarker levels. (**G**) Cox proportional hazards modelling time to MI incidence following MI stratified by diabetes diagnosis. (**H-I**) Effects of known MI biomarkers on time to MI incidence. Individuals stratified by diabetes diagnosis and further into the high quantile (top 30%) and bottom quantile (bottom 30%) of the biomarker levels. Cox proportional hazards regression used to determine hazard ratios and statistical significance (E-F, H-I). *P<0.05, **P<0.01, ***P<0.001, ***P<0.0001.

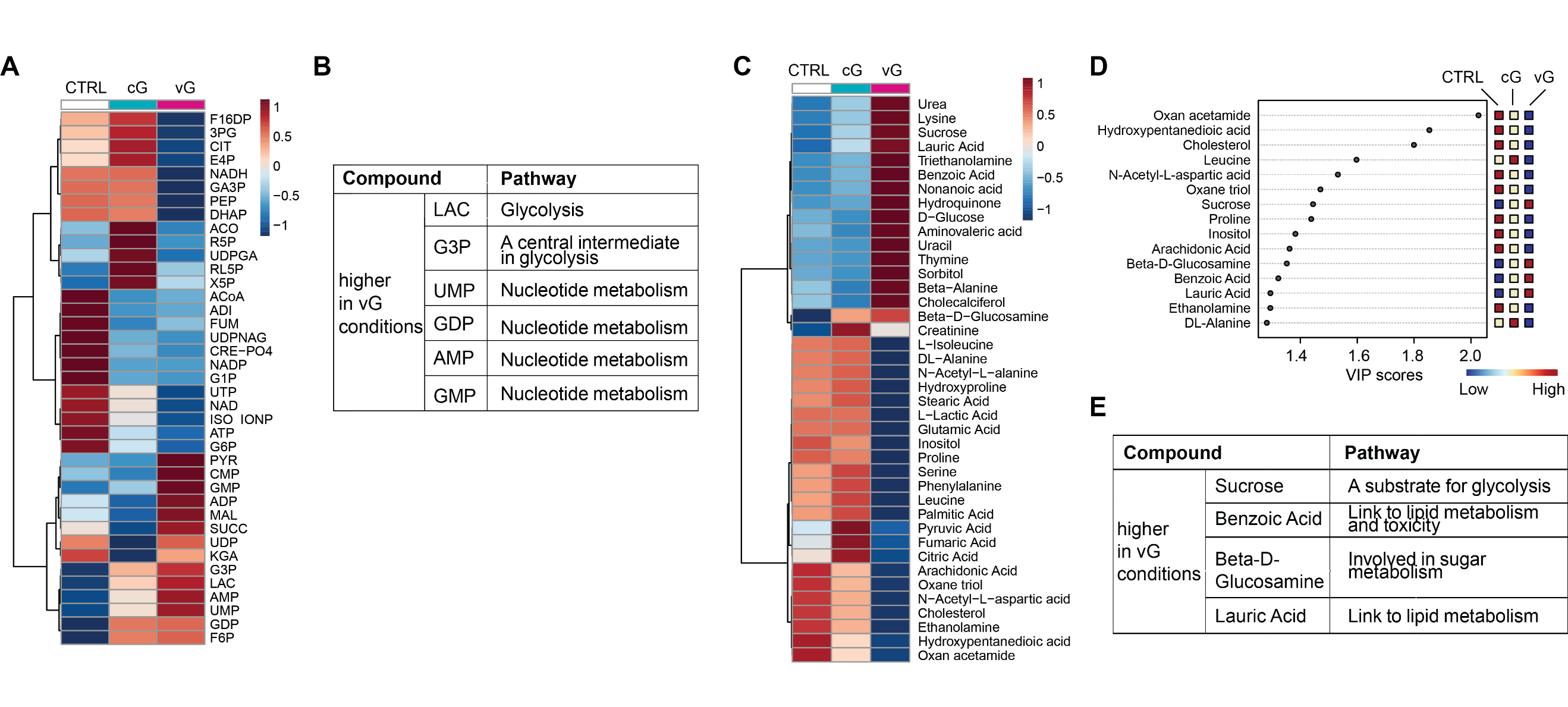
**Supplemental Figure 2. LC-MS and GC-MS metabolomics analysis of control, cG and vG cardiomyocytes.** (**A-B**) LC-MS analysis of the differential concentration of metabolites among control, cG and vG conditions in heatmap (**A**), with a table of higher compounds linked to metabolic pathways (**B**). (**C-E**) GC-MS analysis of the differential concentration of metabolites among control, cG and vG conditions in heatmap (**C**), the relative concentrations of the corresponding metabolite in each group under study (**D**), and a table of higher compounds linked to metabolic pathways (**E**). (n=3; 3 biological replicates each with 2 technical replicates). Statistical analysis by hierarchical clustering analysis (A, C) or partial least squares - discriminant analysis (D).
